# Omitted variable bias in GLMs of neural spiking activity

**DOI:** 10.1101/317511

**Authors:** Ian H. Stevenson

## Abstract

Generalized linear models (GLMs) have a wide range of applications in systems neuroscience describing the encoding of stimulus and behavioral variables as well as the dynamics of single neurons. However, in any given experiment, many variables that impact neural activity are not observed or not modeled. Here we demonstrate, in both theory and practice, how these omitted variables can result in biased parameter estimates for the effects that are included. In three case studies, we estimate tuning functions for common experiments in motor cortex, hippocampus, and visual cortex. We find that including traditionally omitted variables changes estimates of the original parameters and that modulation originally attributed to one variable is reduced after new variables are included. In GLMs describing single-neuron dynamics, we then demonstrate how post-spike history effects can also be biased by omitted variables. Here we find that omitted variable bias can lead to mistaken conclusions about the stability of single neuron firing. Omitted variable bias can appear in any model with confounders – where omitted variables modulate neural activity and the effects of the omitted variables covary with the included effects. Understanding how and to what extent omitted variable bias affects parameter estimates is likely to be important for interpreting the parameters and predictions of many neural encoding models.

## Introduction

Regression models have been widely used in systems neuroscience to explain how external stimulus and task variables as well as internal state variables may relate to observed neural activity (Brown et al., 2003; Kass et al., 2005). However, in many cases, the full set of variables that explain the activity of the observed neurons is not observed or is not even known. It is important to recognize that, in these cases, omitted variables can cause the parameter estimates for the effects that are included in a regression model to be biased. That is, parameter estimates for the modeled effects would be different if other, omitted variables were to be included in the model (Box, 1966). In experiments from behaving animals (Niell and Stryker, 2010; Reimer et al., 2014), but also in more controlled sensory tasks (Kelly et al., 2010; Arandia-Romero et al., 2016), there is growing evidence that neural activity is affected by many more variables than are typically considered relevant (Kandler et al., 2017; Stringer et al., 2018). At the same time, although it has long been a concern in statistics (Pearl, 2009) and has received some attention in other fields (Greenland, 1989; Clarke, 2005), omitted variable bias, as a general problem, appears underappreciated in systems neuroscience. Here we demonstrate why systematically considering omitted variable bias may be important in neural data analysis and examine how omitted variable bias can affect one popular framework for describing neural spiking activity – the generalized linear model (GLM) with Poisson observations.

In general, regression methods aim to estimate variations in a response variable as a function of other variables or covariates. When the goal of modeling is to maximize prediction accuracy, such as with brain machine interfaces, interpreting the model parameters may not be a high priority. However, in many other cases, parameter estimates are, at least to some extent, interpreted and analyzed. For instance, tuning curves or receptive fields may be measured and compared under different stimulus or task conditions or before and after a manipulation. In fully controlled experiments where the covariates are assigned at random, estimated coefficients can often be interpreted as estimates of causal effects (Gelman and Hill, 2007). However, for many cases in neuroscience, it may be difficult or impossible to completely control or randomize all the relevant variables.

In modeling neural activity, omitted variable bias can appear in any situation where neurons are modulated by omitted variables and the omitted variables (often called confounders) are not independent from the variables included in the model – the ones whose effects we are trying to estimate. Minimizing the influence of confounding variables is a major part of most experimental design (Rust and Movshon, 2005), and the statistical effects of confounding variables are well understood (Wasserman, 2004). However, when the goal of modeling is description or explanation (Shmueli, 2010), the effects of these omitted variables are frequently neglected. To give a concrete example, imagine an idealized neuron in primary motor cortex (M1) whose firing, unlike typical M1 neurons (Georgopoulos et al., 1982), is not at all modulated by reach direction but, instead, is modulated by reach speed (Fig 1). In a typical experimental setting, an animal’s reach directions are randomized, but reach speed cannot be randomized or tightly controlled. If the average speed differs across reach directions, such a hypothetical neuron will appear to be tuned to reach direction, despite not being directly affected by direction. First, fitting a typical tuning curve for reach direction, we would infer that such a neuron has a clear preferred direction and non-zero modulation depth. On the other hand, if we then fit a second model that included both reach direction and speed, we would infer that the neuron is modulated by speed alone, and it would be apparent that the original preferred direction and modulation depth estimates were biased due to the omitted variable.

**Figure 1:**
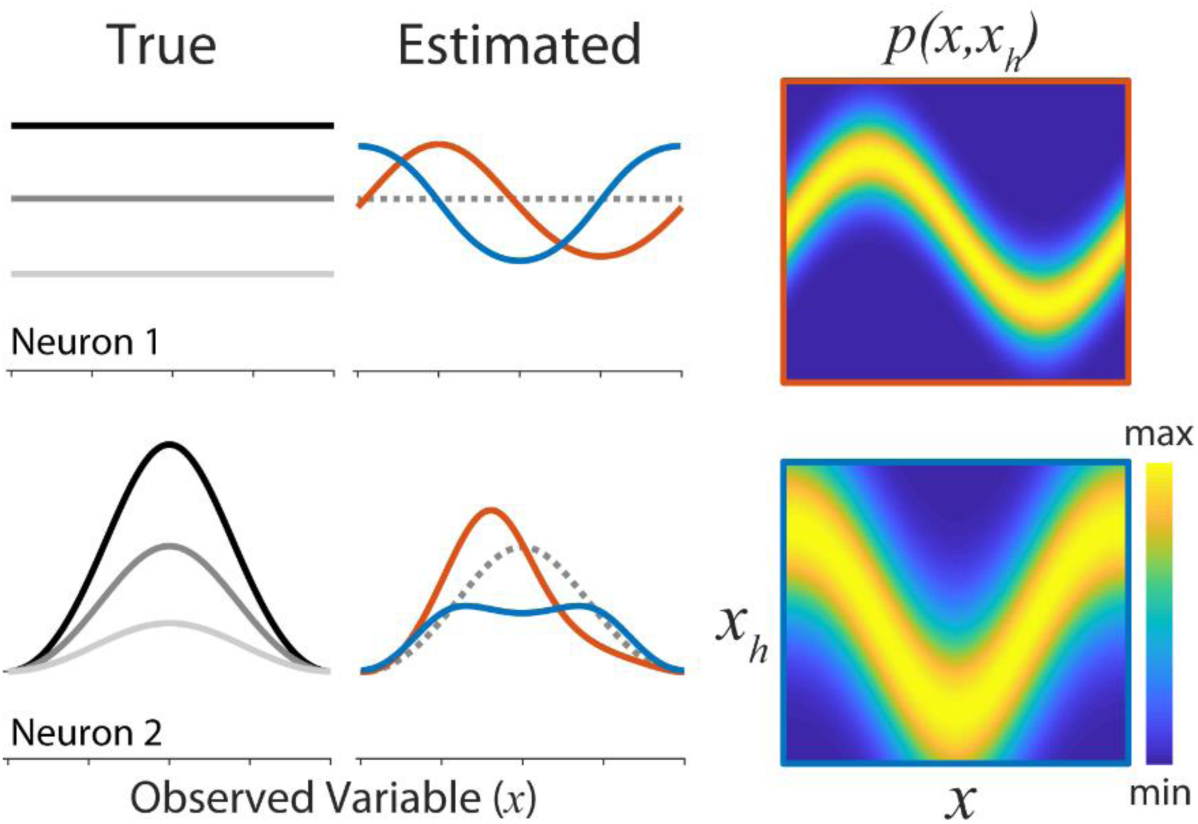
When relevant variables are omitted from the model, estimates of the included effects can be biased. Consider two hypothetical neurons tuned to an observed variable *x* and an omitted variable *x*_*h*_. Neuron 1 is not tuned to the observed variable *x*, but its rate is modulated by the omitted variable with true tuning curves denoted by the gray curves (top left). If *x* and *x*_*h*_ covary, the apparent tuning of this neuron to *x* when the tuning curve is estimated using *x* alone will then be biased (red and blue curves, top middle). This neuron will appear to be tuned to x despite not actually being tuned to this variable. In addition, to this type of illusory tuning, there can also be more subtle biases. Here, neuron 2 is tuned to *x* and *x*_*h*_ (gray curves, bottom left). However, depending on how *x* and *x*_*h*_ covary, the preferred stimulus or the modulation can be misestimated. Here the true tuning to *x* is shown at three different fixed values of *x*_*h*_ (three gray curves, left panels). The estimated tuning when *x*_*h*_ is omitted are shown at center with red and blue curves corresponding to the estimates under two different joint distributions (matching borders, right). Dashed line denotes the effect of *x* when *x*_*h*_ is fixed.

In adding additional variables, previous studies have largely focused on the fact that including previously omitted variables improves model accuracy or the fact that neural activity is often influenced by a host of task variables. In M1, for instance, including speed improves model accuracy (Moran and Schwartz, 1999), but the presence of many correlated motor variables (e.g. kinematics, end-point forces, muscle activity) makes it difficult to interpret how neurons represent movement overall (Humphrey et al., 1970; Omrani et al., 2017). Here, instead of focusing on the advantages or complexities of models with many variables, we focus on the well-known, but under-discussed, fact that the parameters describing the original effects change as additional variables are included. The hypothetical M1 neuron above points to a more general question about regression models of neural activity. What happens when we cannot or do not include variables that are relevant to the process that we are modeling?

Here we first evaluate the statistical problem of omitted variable bias in the canonical generalized linear model with Poisson observations. Then, as a case study, we examine how speed affects estimates of direction tuning of neurons in primary motor cortex, as well as, two other case studies where the spike counts are modeled as a function of external variables: orientation tuning in primary visual cortex (V1) and place tuning in the hippocampus (HC). In each of these case studies we find that commonly omitted variables (speed in M1, population activity in V1, and speed and heading in HC) can bias the estimated effects of commonly included variables (reach direction in M1, stimulus orientation/direction in V1, and place in HC). Across all three case studies, including the omitted variables reduces the estimated modulation due to typical tuning effects. We also illustrate how omitted variable bias can affect generalized linear models of spike dynamics where a post-spike history filter aims to describe refractoriness and bursting (Truccolo et al., 2005). The goal of these models is typically to differentiate aspects of spike dynamics that are due to the neurons own properties (e.g. membrane time constant, resting potential, after-hyper-polarization currents) from those due to input to the neuron from other sources (Brillinger and Segundo, 1979; Paninski, 2004). In this setting, the input to the neuron is typically not directly observed, but is approximated by stimulus or behavioral covariates, local field potential, or the activity of other neurons. Here we show that omitting the input can lead to large biases in post-spike history filters, and that including omitted variables describing the input can change the interpretation and stability of the estimated history effects.

GLMs have been used in many settings to disentangle the effects of multiple, possibly correlated, stimulus or task variables (Fernandes et al., 2014; Park et al., 2014; Runyan et al., 2017) and also to model neural mechanisms such as post-spike dynamics, interactions between neurons, and coupling to local fields (Harris et al., 2003; Truccolo et al., 2005; Pillow et al., 2008). It is often argued that GLMs are advantageous because they have unique maximum likelihood estimates and can be more robust to non-spherical covariate distributions than other methods, such as spike-triggered averaging (Paninski, 2004; Pillow, 2007). Although these advantages are important, GLMs are not immune to bias. Here we show how the possibility of omitted variable bias, in particular, should encourage researchers to be cautious in their interpretation of model parameters, even in cases where a GLM achieves high predictive accuracy (Shmueli, 2010).

## Results

Here we introduce the problem of omitted variable bias and examine differences between omitted variable bias in linear models and the canonical Poisson GLM. We then consider three tuning curve estimation problems: estimating direction tuning in primary motor cortex, place tuning in hippocampus, and orientation tuning in primary visual cortex and show how omitted variables in each of these three cases can alter parameter estimates. Finally, we consider a GLM that aims to describe the dynamics of post-spike history and show how omitted inputs can bias the estimated history effects and qualitatively change model stability.

### Omitted Variable Bias in Linear Regression and canonical Poisson GLMs

When relevant variables are not included in a regression model, the estimated effects for the variables that are included can be biased (Box, 1966). Omitted variable biases can cause the parameters describing the effects of the original variables to be over- or under-estimated, and model fits can change qualitatively when omitted variables are included (Fig 1).

To understand the problem of omitted variable bias it will be helpful to briefly review the well-known case of multiple linear regression, where the bias can be described analytically (Box, 1966). In the linear setting, consider the generative model

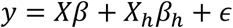

where observations *y* are a linear combination of observed *X* and omitted *X*_*h*_ variables plus normally-distributed i.i.d. noise *ϵ∼N*(0, *σ*). For simplicity, we ignore the intercept term, but in the analysis that follows it may also be considered as part of *X*. If we then fit the (mis-specified) model without *X*_*h*_ using maximum likelihood estimation (equivalent to the ordinary least squares solution, in this case) the estimated parameters will be

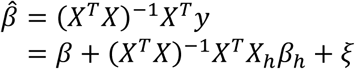

where *ξ* denotes the effect of the noise, and the bias (*X*^*T*^*X*)^−1^*X*^*T*^*X*_*h*_*β*_*h*_ will, generally, be non-zero. There will be no bias only in the cases where the omitted variables do not affect the observations (*β*_*h*_ = 0) or when the omitted variables and observed variables are not collinear (*X*^*T*^*X*_*h*_ = 0). Note that (*X*^*T*^*X*)^−1^*X*^*T*^*X*_*h*_ is the matrix of regression coefficients for the omitted variables using the observed variables as predictors. For linear regression, the omitted variable bias thus depends on both the extent to which the omitted variables affect the observations *β*_*h*_ and the extent to which the omitted variables can be (linearly) predicted from the observed variables.

Although there is a closed-form solution for the omitted variable bias for linear regression, the generalized linear setting is not as tractable (Gail et al., 1984; Clogg et al., 1992; Drake and McQuarrie, 1995). We will consider the case of a canonical Poisson GLM, in particular, where

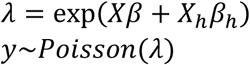

In the more general case, GLMs have

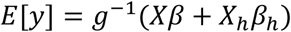

where *g*^−1^(·) is the inverse link function, and *y* is distributed following an exponential family distribution (McCullagh and Nelder, 1989). For a canonical GLM the log-likelihood takes the form

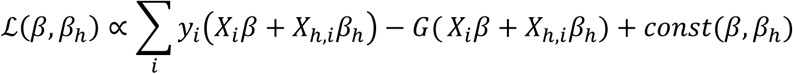

where the nonlinear function *G*(·) depends on both the link function and the noise model. For canonical GLMs, this log-likelihood is concave and the maximum likelihood estimate 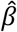 satisfies 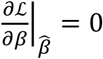. The exact form of *G*(·) will depend on the model, but for linear regression *G*(*x*) is proportional to 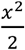, and for canonical (log-link) Poisson regression *G*(·) = exp(·).

Now, with omitted variables, instead of maximizing the correct log-likelihood, we maximize instead

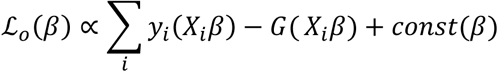

For the omitted variable bias in 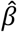 to be 0, we need both 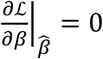 and 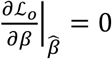 at the same value of *β*. Although, neither MLE has a closed form solution, this condition implies that, if there is no bias due to the omitted variables,

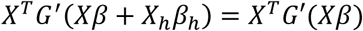

where *G*^′^(·) is the derivative of *G*(·). For linear regression this equality reduces to the OLS form derived above, and for canonical Poisson regression we have

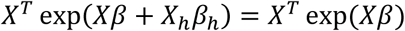

This equality is satisfied when observations are not modulated by the omitted variables *β*_*h*_ = 0 or, more generally, when the effect of the omitted variables δλ = exp(*Xβ* + *X*_*h*_*β*_*h*_) – exp(*Xβ*) is orthogonal to the included variables *X*. Note that with linear regression, *X*^*T*^*X*_*h*_ = 0 implies that the estimates will not be biased, but here this is not the case unless *X*^*T*^*δλ* = 0 as well. Due to the structure of the canonical Poisson GLM, omitted variable bias can thus occur even in a properly randomized, controlled experiment (Gail et al., 1984),

It is important to note that the maximum likelihood estimates themselves are consistent. That is, the estimators converge (in probability) to their true values when the generative model is correct. The bias here is a result of the model being mis-specified. This mis-specification affects the location of the maximum and, also, the shape of the likelihood. Optimization methods, such as Newton’s method, will typically contain omitted variable bias in each parameter update. For canonical Poisson regression, for instance, the updates take the form

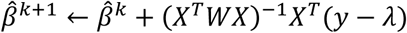

at iteration *k* where the weight matrix *W* is diagonal with entries *W*_*ii*_ = *λ*_*i*_ and (*X*^*T*^*WX*)^−1^ is the Fisher scoring matrix (inverse Hessian of the log-likelihood) at the current estimate 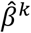. Since the mis-specified model will use *λ* = exp(*Xβ*) instead of the exp(*Xβ* + *X*_*h*_*β*_*h*_), both the weight matrix and the gradient *X*^*T*^(*y* – *λ*) will be biased at each step of the optimization (except when 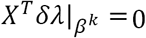). Traditional standard errors for the MLE will also typically be influenced by omitted variables, since 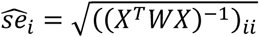. Moreover, as previous studies have shown, omitted variables can lead to misestimation of the variability in *E*[*y*] and dispersion *var*(*y*) (Czanner et al., 2008; Goris et al., 2014). If the omitted variables affect the observations, then they will generally increase the variability of *E*[*y*]. Then, unless the omitted variables are perfectly predicted by the included variables, the explained variance *var*(*E*[*y*]) of the mis-specified model will be lower than that of the full model. This may, in turn, lead to overestimates of dispersion, since *var*(*y*) = *E*[*var*(*y*)] + *var*(*E*[*y*]).

### Omitted Variable Bias in Tuning Curve Estimation

When fitting tuning curve models to spike count data, omitted variable bias can cause preferred stimuli and modulation depths to be misestimated and can even lead to completely illusory tuning (Fig 1). To illustrate how omitted variable bias affects GLMs of neural spiking, not just in theory, but in practice, we consider three case studies where we fit typical tuning curve models that omit potentially relevant variables along with augmented models that include these additional variables. We first consider modeling spike counts across trials and on slow (>100ms) timescales. Here we assess 1) the tuning of neurons in motor cortex to reach direction, with speed as a potential omitted variable, 2) the tuning of neurons in hippocampus to position, with both speed and head-direction as potential omitted variables, and 3) the tuning of neurons in visual cortex to the direction of motion of a sine-wave grating, with population activity as a potential omitted variable. In each of these cases studies, we show how the omitted variables are not independent from the commonly included variables and how neural responses are modulated by the omitted variables. These two properties, together, can lead to omitted variable biases.

In our first case study, we model data recorded from primary motor cortex (M1) of a macaque monkey performing a center-out, planar reaching task. In this task, speed differs systematically across reach directions (Fig 2A), with average speed differing by as much as 35±3% (for the targets at 45 and 225 deg relative to right, Fig 2B). To model neural responses, we first fit a traditional tuning curve model (Georgopoulos et al., 1982; Amirikian and Georgopulos, 2000), where the predicted responses depend only on target direction. Here we use a circular, cubic B-spline basis (5 equally spaced knots) to allow for deviations from sinusoidal firing, but, in most cases, the responses of the n=81 neurons in this experiment are well described by cosine-like tuning curves with clear modulation for reach direction. We then fit a second model that includes effects from movement speed. Here we use covariates based on (Moran and Schwartz, 1999), including a linear speed effect, as well as, cosine-tuned direction-speed interactions (see Methods). This model captures the responses of individual neurons, where spike counts can increase (Fig 2C, top) or decrease (Fig 2C, middle) as a function of speed, and, in some cases, speed and direction appear to interact (Fig 2C, bottom). Together, the fact that direction and speed are not independent along with the fact that neural responses appear to be modulated by speed could lead to biased parameters estimates for the model where speed is omitted.

Comparing the models with and without omitted variables we find that, averaged across the population, there are only minimal shifts in the preferred direction (3±2 deg) when speed is included in the traditional tuning curve model, and there do not appear to large, systematic shifts in the population distribution of PDs (Kuiper’s test, p>0.1). At the same time, there is substantial variability between neurons in the size of the PD-shift (circular SD 32±5 deg). Across the population, modulation depth (measured using the standard deviation of the tuning curve) decreases slightly on average (3±2%), and the size of the modulation change also varies substantially between individual neurons (SD of changes 18±3%). An example neuron in Fig 1C (bottom), for instance, has a modulation decrease of 9±5% and the preferred direction changes 4±9 deg when speed is included in the model (standard error from bootstrapping). Overall, ∼10% of neurons have statistically significant changes in PD, and ∼14% have significant changes in modulation (bootstrap tests *α*=0.05, not corrected for multiple comparisons). For some individual neurons, at least, the parameters of the model without speed, thus, have clear omitted variable bias. However, since individual neurons have diverse speed dependencies, in this case, the average biases across the population are minimal.

When speed is included in the model, model accuracy (pseudo-R^2^) does increase slightly (p=0.01, one-sided paired t-test across neurons). The average cross-validated (jack-knife) pseudo-R^2^ for the original model is 0.23±0.01 and for the model with speed 0.24±.01 (Fig 5). However, it seems likely that in other experimental contexts the effects of omitting speed could be more pronounced. By requiring the animal to make reaches to the same targets at different speeds, previous studies have more clearly demonstrated that responses in M1 are modulated by speed (Churchland et al., 2006). Here we demonstrate how this type of modulation can lead to omitted variable biases in the estimated parameters of typical tuning curve models without speed.

**Figure 2:**
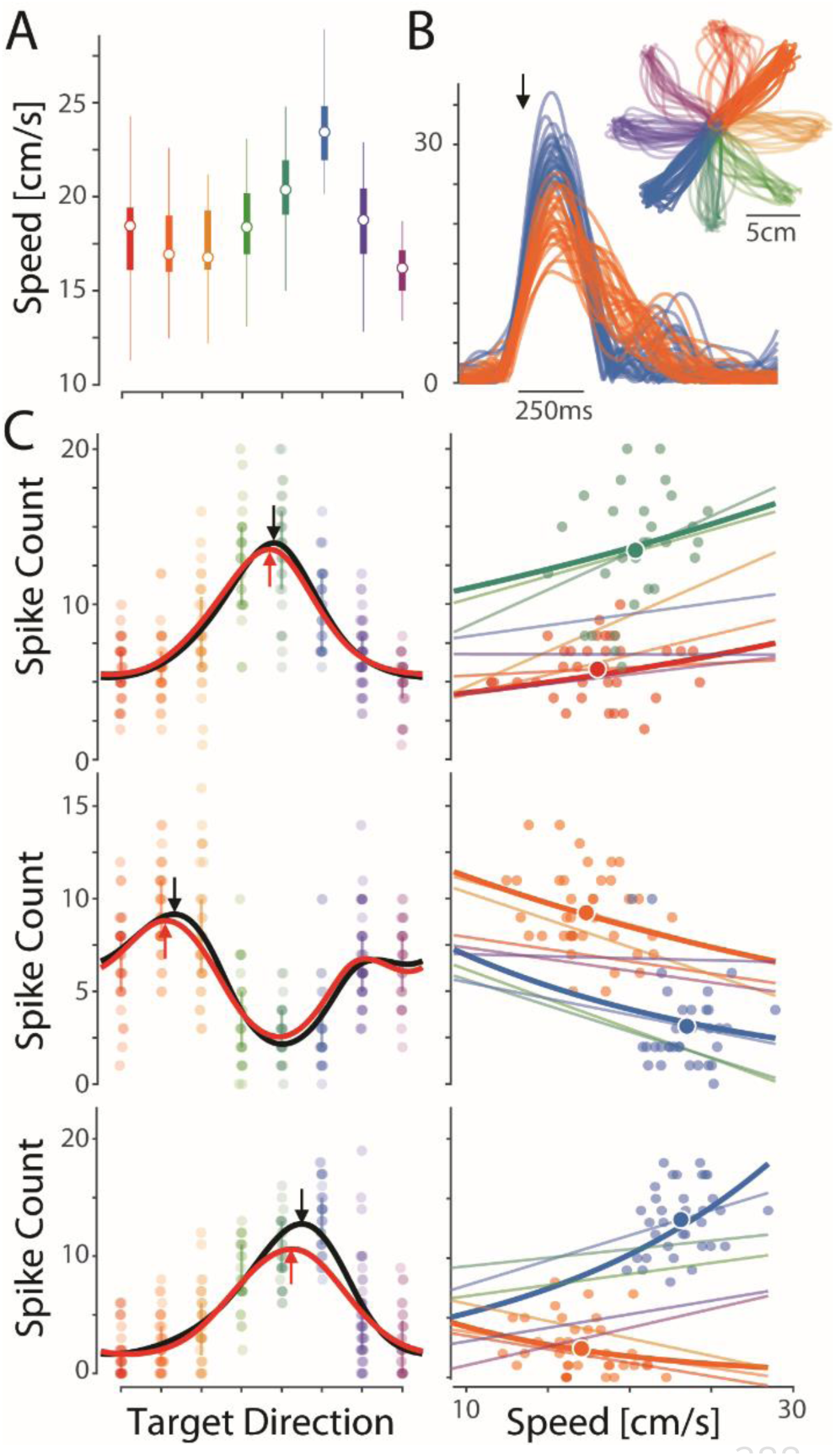
Speed as an omitted variable in M1 tuning for reach direction. A) The distribution of reach speeds differs by target direction in a center-out task. Circles denote median, boxes denote IQR. B) Speed profiles for the two targets showing the largest speed differences. Individual traces denote individual trials aligned to the half-max (black arrow). Inset shows the position of each trial with colors denoting reach direction. C) The responses of 3 M1 neurons show typical tuning for reach direction. The tuning curve estimated using direction covariates alone (black) changes when speed covariates are included (red). Red curves denote the direction effect within the full model and are generated by assuming speed is constant (equal to the mean speed across all trials). Right panels illustrate the speed dependence for the preferred direction and its opposite. Dark lines denote the estimated effect of speed under the full model. Data points show single trial data, along with the mean speed and rate for each direction (big data point). Light lines show linear trends (OLS) using only the trials from each specific target.

In our second case study we examine the activity of neurons in the dorsal hippocampus of a rat foraging in an open field. Here we consider to what extent the practice of omitting speed and head direction from a place field model biases estimates of a neuron’s position tuning. As in the first case study, omitted variable bias can occur if neural activity is modulated by omitted variables and the omitted variables covary with the included variables. In the case of the hippocampus, neural activity is known to be modulated by both movement speed and head direction (McNaughton et al., 1983), in addition to an animal’s position (O’Keefe and Dostrovsky, 1971). Additionally, behavioral variables can be highly nonuniform across the open field (Walsh and Cummins, 1976), for instance, near and far from the walls. Together the fact that the omitted variables may covary with position and the fact that neurons appears to be modulated by the omitted variables, suggest that there may be omitted variable bias.

Here, in one recording during an open field foraging task we find that the average speed (Fig 2A) and heading (Fig 2B) differ extensively as a function of position. Within a given neuron’s place field, the distributions of speed and heading may be very different from their marginal distributions. Across the population of n=68 place cells (selected from 117 simultaneously recorded neurons, see Methods), average in-field speed was between 80-135% of the average overall speed (5.5cm/s), and the animal’s heading can be either more or less variable in-field (circular SD 57-80 deg) compared to overall (75 deg).

As previous studies have shown, we also find that neural responses are modulated by speed and head direction. Responses due to place, speed, and heading are shown for one example neuron in Fig 3. This neuron shows a stereotypical place-dependent response (Fig 2B), but splitting the observations by speed (Fig 3C, top) or heading (Fig 3B, bottom) by quartiles/quadrants reveals that there is also tuning to these variables. The neuron appears to increase its firing with increasing speed and responds most strongly when the rat is facing the left. These dependencies are well fit by the full model where the firing rate depends, not just on position, but also on the (log-transformed) speed and the heading (Fig 3D, bottom). For the example place cell shown here, the location of the place-field does not change substantially when the omitted variables are included (Fig 3E). However, the modulation (SD of the rate map) decreases by 27%. That is, 27% of the apparent modulation due to position when it is modeled alone, can be explained by speed and heading effects.

Across the population of place cells, there were no clear, systematic difference in the place field locations, but the modulation (SD of the rate map *λ*(*x*)) decreases by 9±1% on average when speed and heading are included. Individual neurons showed substantial variability in their modulation differences (population SD 10±1%). As in M1, including the omitted variables increased spike prediction accuracy – the average cross-validated (10-fold) pseudo-R^2^ was .29±.02 for the original model and .31±.02 for the model including speed and heading activity. This difference seems small, since there is large variability in pseudo-R^2^ values across the population, but the average increase in pseudo-R^2^ was 11±3% (Fig 5). Given that neurons appear to be modulated by speed and heading, it is unsurprising that including these variables improves model fit. However, as before, it is important to note that this modulation can lead to biases in the place field estimates for the model with only position.

**Figure 3:**
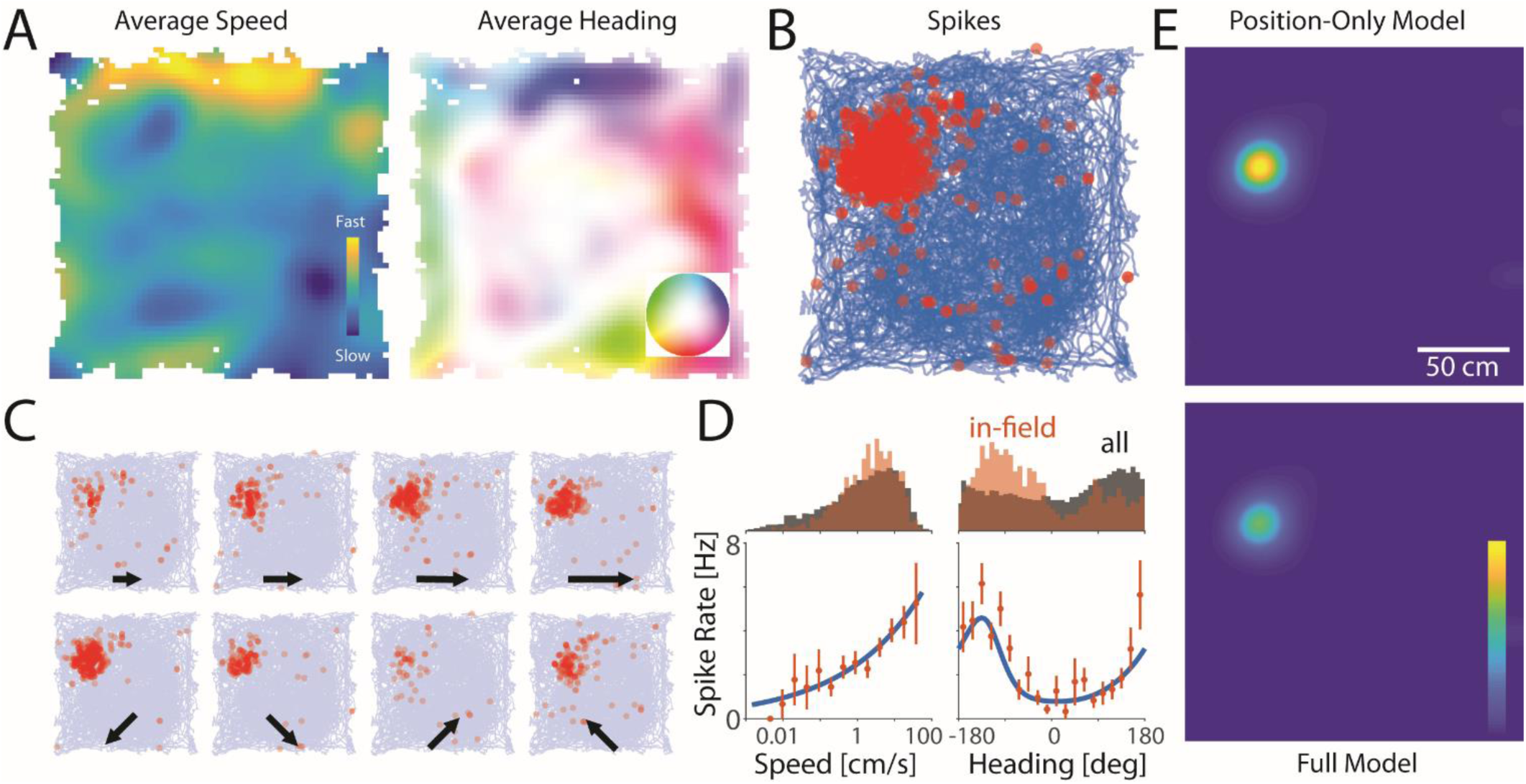
Speed and heading as omitted variables in hippocampal place cells. A) Average speed and heading as a function of position for a rat foraging in an open field. B) An example place cell tends to spike (red dots) when the animal is at a specific position in space. C) The activity of this neuron is modulated by the animal’s speed (top row) and heading (bottom row). Speed is split into quartiles, subplots include all headings. Heading is split into quadrants, subplots include all speeds. D) The distributions of speed and heading within the place field differ from the overall distributions, and the neuron is tuned to these variables. Blue curve shows model fit. E) After modeling the effect of speed and heading within the place field, the location of the place field does not change but the apparent modulation due to position is reduced.

In our third case study, we examine the activity of neurons in a more controlled sensory experiment. Here we use data recorded from primary visual cortex (V1) of an anesthetized monkey viewing oriented sine-wave gratings moving in on of 12 different directions (see Methods). In this experiment, variability in the animal’s behavior is purposefully minimized, and, instead of considering the effect of omitting a behavioral variable, here we consider the effect of omitting a variable relating to the animal’s internal state – the total population activity. Several studies have previously shown that population activity alters neural responses in V1 (Arieli et al., 1996; Kelly et al., 2010; Okun et al., 2015; Arandia-Romero et al., 2016). If the distribution of population activity also varies with stimulus direction, then there is the potential for omitted variable bias.

Here we assess neural activity from n=90 simultaneously recorded neurons across many (2400) repeated trials with 12 different movement directions. We find that there is high trial-to-trial variability in the population rate (Fig 4A), and the average firing across all neurons does differs across stimulus directions, up to ∼50%. For this recording, the most extreme differences were between the 180 deg stimulus where the average rate across the population was 3.4±0.1Hz and the 60 deg stimulus where the average rate was 6.3±0.1Hz (Fig 4B). By adding the (log-transformed) population rate as a covariate to a more typical model of direction tuning, we find that population activity may lead to omitted variable bias in models of direction tuning alone.

As in the case studies above, there do not appear to be any consistent or systematic effects on the preferred stimulus direction at the population level (Kuiper’s test, p=0.1). However, the modulation depth (measured using SD of the tuning curve) decreases substantially 15±2% when population rate is included in the model, and there is again high variability across neurons (SD 20±2%). In this case, model accuracy increases substantially when the omitted variable is included. The cross-validated (10-fold) pseudo-R^2^ is .26±.02 for the original model and .43±.02 for the model including population activity, with an average increase of 164±31% (Fig 5).

**Figure 4:**
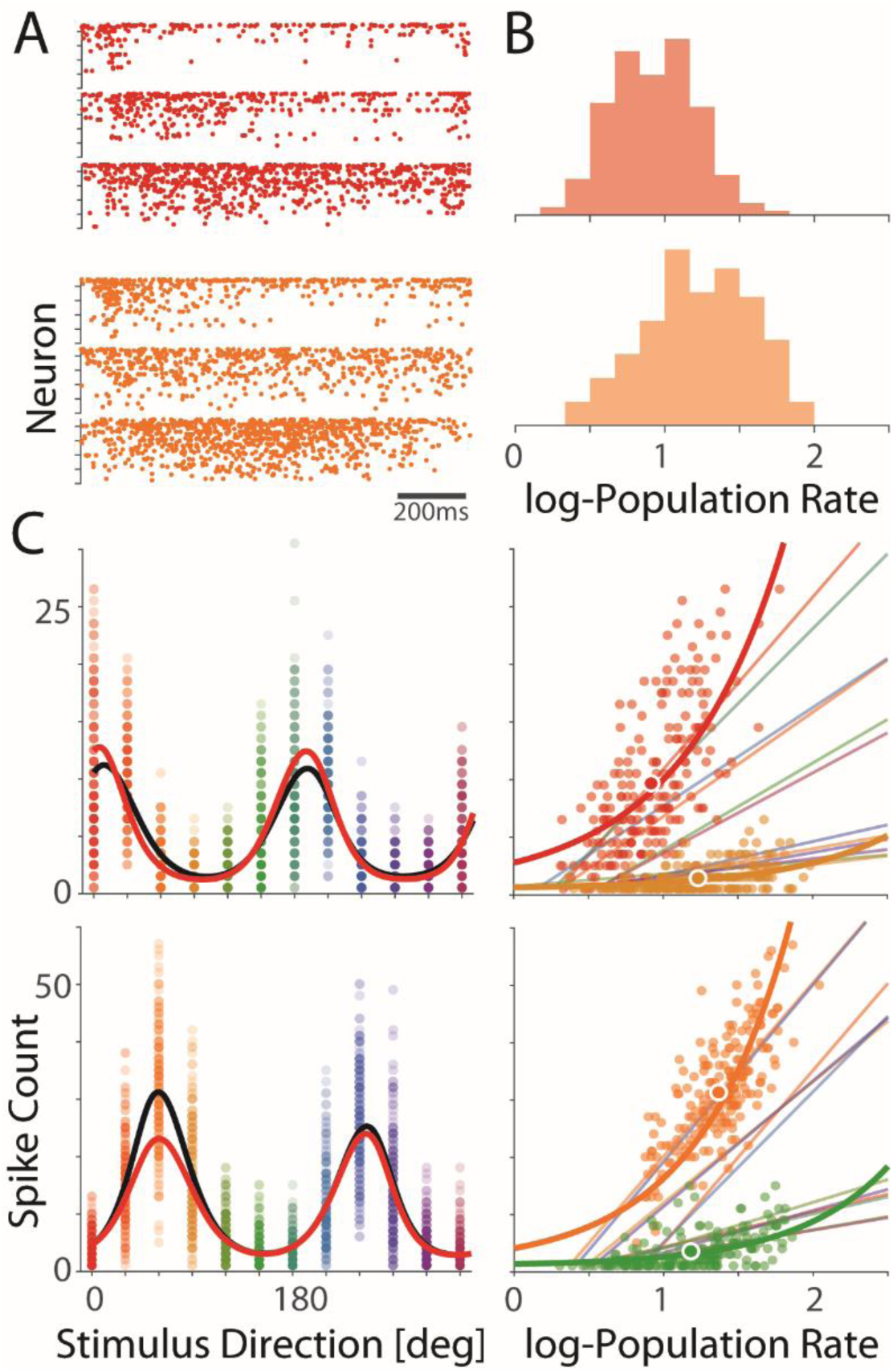
Population rate as an omitted variable in primary visual cortex. A) Correlated trial to trial variability. Population rasters for three trials of the same drifting grating stimulus (0 deg, red and 30 deg, orange). Neurons are sorted by overall firing rate. B) Histograms of the population rate across trials. As a population, the neurons respond at higher rates to 30 deg stimuli, but there is high trial-to-trial variability. C) The responses of 2 V1 neurons show typical tuning for direction of motion. The tuning curve estimated using direction covariates alone (black) changes when the population rate covariate are included (red). Right panels illustrate the dependence for the preferred direction and an orthogonal direction. Dark lines denote the estimated effect of speed under the full model. Data points show single trial data, along with the mean count and rate (big data point). Light lines show linear trends (OLS) using only the trials from each specific stimulus.

Unlike in M1 where the effect of speed was highly diverse for different neurons, in this case study the effect of the population rate is largely consistent. Higher population rates are associated with higher firing rates, and, for most neurons, the effect of the population rate is stronger in the preferred direction(s), consistent with a multiplicative effect. Note that here, we do not include the neuron whose rate we are modeling in the calculation of the population rate. However, using the population rate as an omitted variable requires some interpretation. The population rate will certainly be affected by the tuning of the, relatively small, sample of neurons that we observe. If we have a disproportionate number of neurons tuned to a specific preferred direction, the population rate in those directions will be higher. This suggests that in a different recording, the covariation between the stimulus and the population rate could very likely be different. However, it appears that the omitted variable biases in this case are mostly driven by noise correlations, where neural activity is correlated on single trials even within the same stimulus condition, rather than stimulus correlations, where neural activity is correlated due to similar tuning. When we shuffle the data within each stimulus condition (removing noise correlations) the average change in the modulation depth is -1±2% (SD 18±3%), and the effect of the omitted variable becomes negligible.

**Figure 5:**
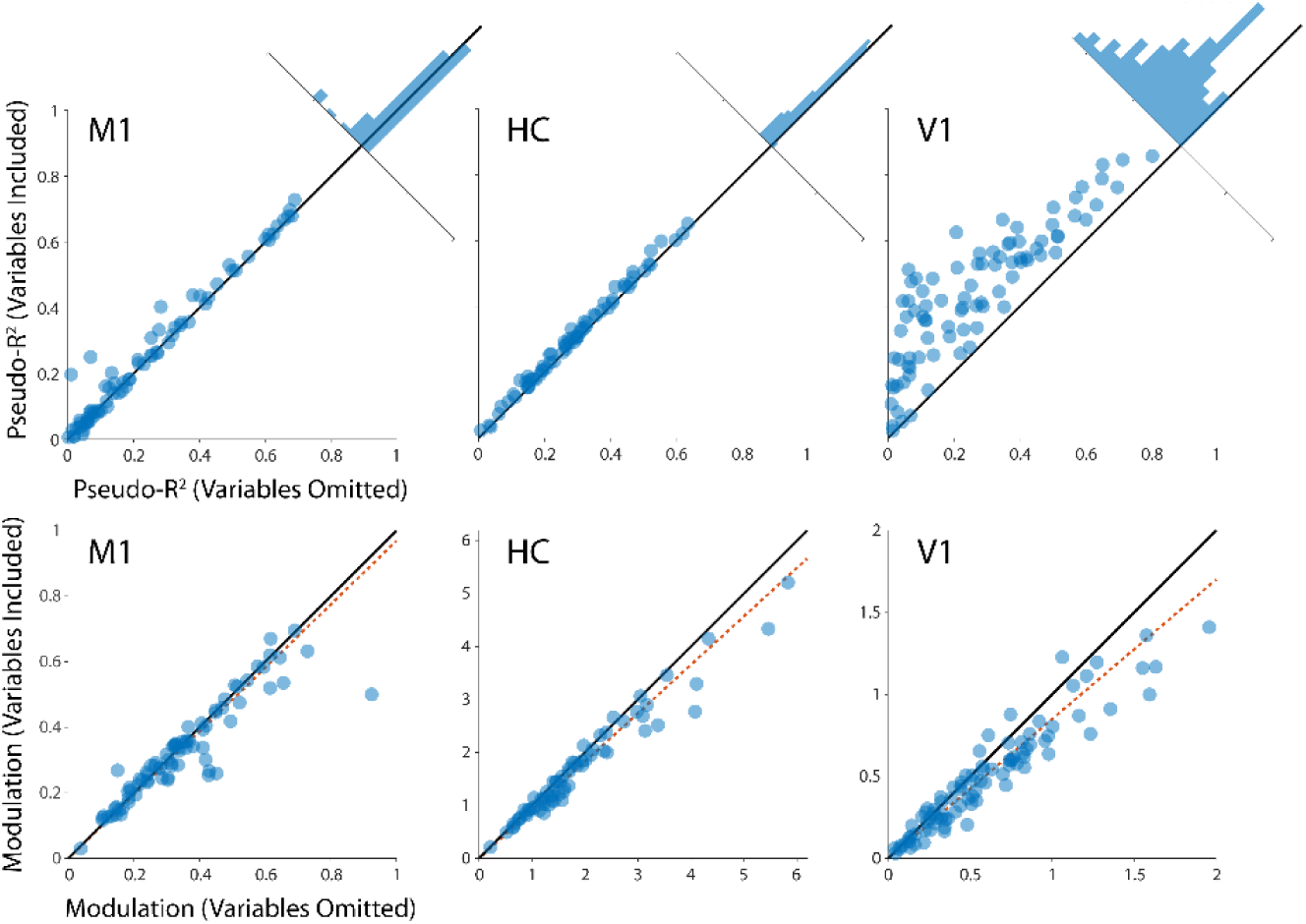
For each of the case studies, on average, the model accuracy increases when omitted variables are included (top) and the modulation due to the original variables decreases (bottom). Scatter plots indicate cross-validated pseudo-R2 values for each neuron under the two models. Modulation denotes the standard deviation of the tuning to the original variable(s) under each model. Here, modulation values are normalized by the average rate of each neuron. Black lines denote equality. Red dashed lines denote linear fit with 0 intercept.

**Figure 6:**
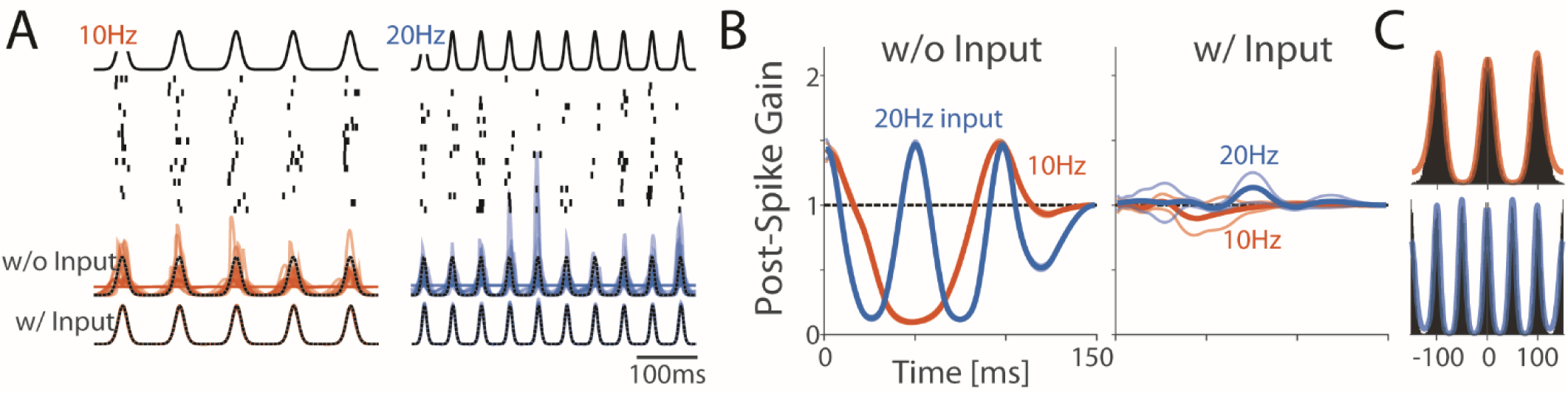
Estimated post-spike history filters can be heavily biased when the input is not included in the model. A) Here we simulate from an inhomogeneous Poisson model with sinusoidal input (no post-spike history effects). The input and spike responses from 20 trials are shown. Although there are no history effects in the generative model, a GLM with history effects that is missing the correct input covariate will use the history terms to capture the structure in the autocorrelation (C). Traces denote the estimated rate for the 20 trials shown above. When the history term is included in the model, but the input is not, the GLM can still reconstruct PSTH responses using the post-spike history alone. B) Post-spike filters for the models in (A) with 95% confidence bands. Note that when input is included in the model the filters correctly reconstruct the true (lack of) filter, and that there is higher uncertainty around the regions where the ISI distribution does not constrain the model.

### Omitted Variable Bias in the Estimation of Post-Spike History Effects

In addition to modeling spike counts over trials or on relatively slow (>100ms) behavioral timescales, GLMs are also often used to describe detailed, single-trial spike dynamics on fast (<10ms) timescales. One common covariate used in these types of models is a post-spike history effect where the probably of spiking at a given time depends on the recent history of spiking. Modeling these effects allows us to describe refractoriness, bursting (Paninski, 2004; Truccolo et al., 2005), and a whole host of other dynamics (Weber and Pillow, 2017). Conceptually, the goal of these models is to disentangle the sources of rate variation based only on observations of a neuron’s spiking, with history effects, ideally, reflecting intrinsic biophysics. However, since the full synaptic input is typically not known with extracellular spike recordings there is potential for omitted variable biases.

To illustrate the potential pitfalls of omitting the input to a neuron, consider using the GLM to capture single neuron dynamics in the complete absence of external covariates

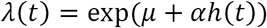

where the rate *λ* is determined by a baseline parameter *μ* along with a filtered version of the neuron’s past spiking with *h*_*i*_(*t*) = ∑_τ>0_ *f*_*i*_(τ)*n*(*t* – τ). This is a perfectly acceptable model of intrinsic dynamics, but for most spike data that we observe this isolated neuron model may not provide a realistic description of a neuron receiving thousands of time-varying synaptic inputs. If we fit this model to data where the input to the neuron did vary over time,

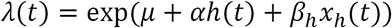

then the history filter in the first model will attempt to capture variation in spiking due to the time-varying input, in addition to any intrinsic dynamics. For example, when *x*_*h*_ is periodic, the estimated history filters of the original model will attempt to capture this periodic structure (Fig 6A-B). Just as in the tuning curve examples above, the fact that history effects covary with the input and the fact that the input modulates the neuron’s firing leads to omitted variable bias. When the input is omitted from the model, the biased history effects simply provide the best (maximum likelihood) explanation of the observed spiking (Fig 6C).

These examples with strong, periodic input are not necessarily biologically realistic, but they make it apparent how the post-spike history can be biased by omitted input variables. In vivo, neurons instead appear to be in a high-conductance state, where membrane potential fluctuations have approximately 1/*f* power spectra (Destexhe et al., 2001, 2003). When these naturalistic input statistics are used to drive the GLM, omitted variable bias can occur, as well. Here we simulate a GLM receiving 1/*f*^*α*^ noise input with *α* = 0 (white noise) 1 and 2 (Fig 7). For white noise input, the MLE accurately recovers the simulated post-spike history filter when the input is omitted from the model, but when *α* = 1 or 2 the estimates become increasingly biased (Fig 7A,C). With the full model, where the input is included as a covariate, the history is recovered accurately no matter what the input statistics are. Just as in the periodic case, however, these different input statistics alter the auto-correlation, and, when the input is omitted from the model, the maximum likelihood history filter simply aims to capture these patterns.

**Figure 7:**
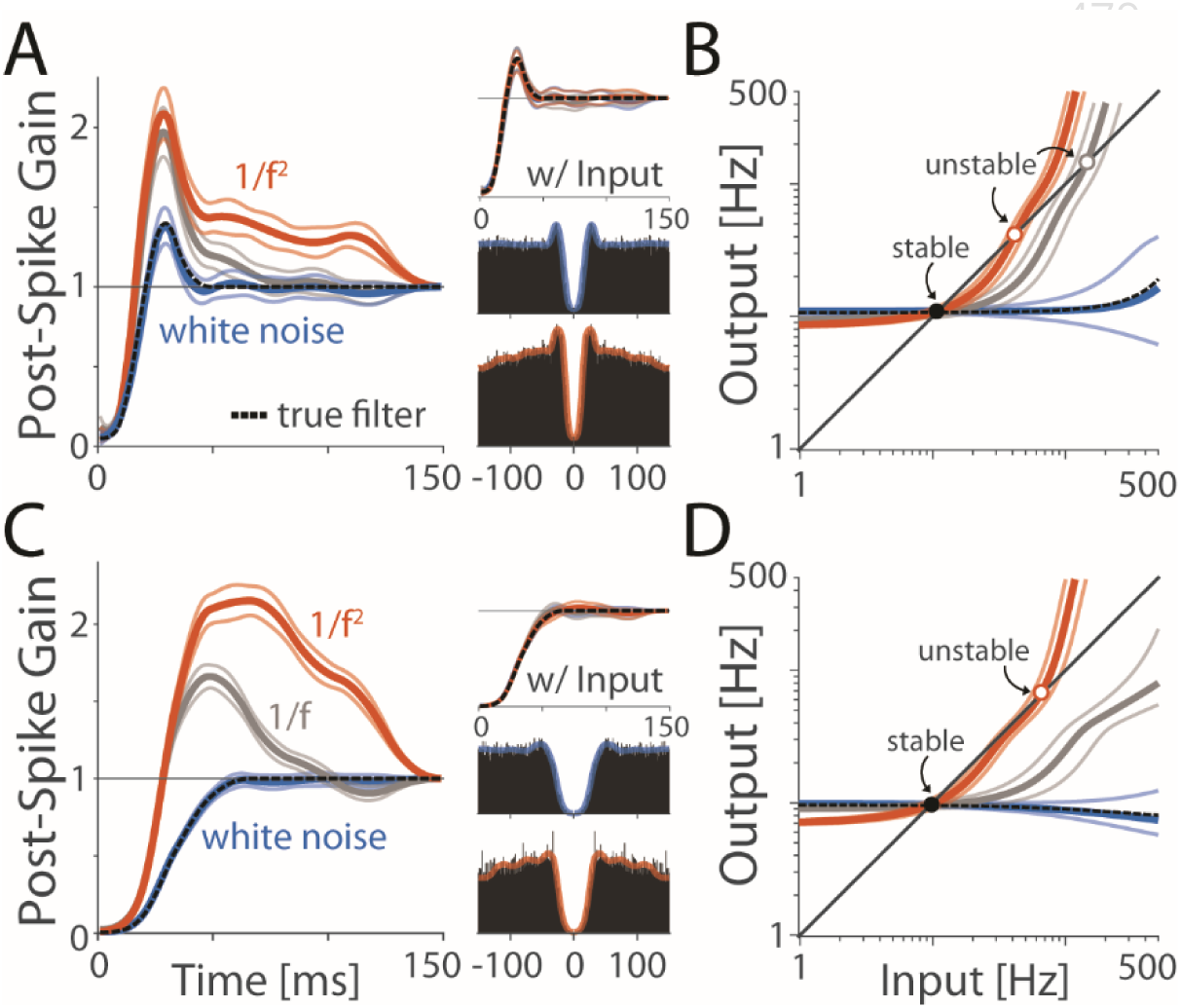
Post-spike filters can show omitted variable bias even in a more realistic scenario. Here we simulate from a GLM with a refractory post-spike filter and drive the neurons with 1/*f*^*α*^ noise. Excepting the case of white noise (*α* = 0), the post-spike filters estimated for the GLM without input are heavily biased (A). C) Even when the effect of the true post-spike filter is to strictly decrease the firing rate, the estimated filters can increase the firing rate. B,D) Approximate transfer functions from a quasi-renewal approximation. When the true filter is stable, the estimated filters can result in fragile dynamics.

In GLMs for single-neuron dynamics, one effect of omitted variable bias is that it may lead us to misinterpret how stable a neuron’s dynamics are. Even if the true history filter only reduces the neuron’s firing rate following a spike (as in Fig 7C), the estimated filter can be biased upwards when the input is omitted. If we were to simulate the activity of this neuron based on the biased filter, the bias could cause the neuron’s rate to diverge if the rate becomes high enough. To assess the stability of the estimated post-spike history effects quantitatively, here we make use of an quasi-renewal approximation analysis introduced in (Gerhard et al., 2017). Given a history filter, this approach finds an approximate transfer function describing the neuron’s future firing rate (output) given its recent (input) firing rate (see Methods). For all estimated models, the transfer function has a stable fixed point near the neuron’s baseline firing rate. When the true input is omitted and *α* > 0, the estimated history filters also have an unstable fixed point where the neuron’s firing rate will diverge if the rate exceeds this point (Gerhard et al., 2017). Here we find that omitted variable bias leads to apparent fragility (Fig 7B,D). The stable region shrinks as *α* increases, and even when the true dynamics are strictly stable (as in Fig 7C,D), omitted variable bias can lead us to mistakenly conclude that the neuron has fragile dynamics.

With most extracellular spike recordings, the synaptic input that the neuron receives is unknown. However, there may also be omitted variable bias when history effects are estimated from real data. In this case, the input to a neuron can be approximated by stimulus or behavioral variables, local field potentials, or the activity of simultaneously recorded neurons (Harris et al., 2003; Truccolo et al., 2005, 2010; Pillow et al., 2008; Kelly et al., 2010; Gerhard et al., 2013; Volgushev et al., 2015). Just as in the simulations above, including or omitting these variables can then alter the estimated history effects, even though they are not as directly related to spiking as the synaptic input itself. Here we consider total population spiking activity as a proxy for synaptic input and consider how including population activity alters the history filters when compared to a model of history alone.

We examine two datasets: spontaneous activity from primary visual cortex of an anesthetized monkey with n=62 simultaneously recorded neurons and activity from dorsal hippocampus of a sleeping rate with n=39 simultaneously recorded neurons. To model population covariates we sum the spiking of all neurons, excepting the one whose spiking we aim to predict, and low-pass filter the signal (see Methods). Similar to previous results (Okun et al., 2015), we find that, since neurons often have correlated fluctuations in their spiking (Fig 8A, D), the population rate is a good predictor for single neuron activity. Moreover, when we add population covariates to a GLM with post-spike history effects the history filter changes.

In the V1 dataset, the post-spike gain decreases by 7.8±0.5% on average when population covariates are included, and 14.9±0.8% when considering only the first 50ms after a spike (Fig 8B). The effects of adding population covariates are less pronounced in the hippocampal dataset. The post-spike gain decreases by 2.5±0.3% on average, and 9.5±1.2% when considering only the first 50ms after the spike (Fig 8E). Based on the quasi-renewal approximation, all neurons in both the V1 and hippocampal datasets have fragile transfer functions where there is a stable fixed point (near the neuron’s average firing rate) and an unstable fixed point where the neuron’s rate diverges if the input becomes too strong. For V1, the average upper-limit of the stable region is 80±3Hz for the models with history only and 143±7Hz for the models with population covariates (Fig 8C). In the hippocampal data, the average upper-limit of the stable region is 38±6Hz for the models with history only and 75±13Hz for the models with population covariates (Fig 8F). Each neuron is, thus, apparently, more stable after the population covariates are included.

As in the case studies using tuning curves, adding covariates also improves spike prediction accuracy. In the V1 dataset, the average log likelihood ratio relative to a homogeneous Poisson model is 2.2±0.3 bits/s for the history model and 3.3±0.3 bits/s for the model with population covariates. In hippocampus, the log likelihood ratio is 0.9±0.3 bits/s for the history model and 2.0±0.5 bits/s for the model with population covariates. The larger effects in V1 are likely explained by the fact that the population rate is predictive for many more neurons here than for the hippocampal data. In the hippocampus, only 26% of the neurons have an increase of over 0.5 bits/s when the population covariates are included, compared to 85% of neurons in V1. Altogether these results demonstrate how omitted variable bias could affect estimates of post-spike history filters in vivo. In both datasets we find that when population covariates are included in the GLM spike prediction accuracy increases, post-spike gain decreases, and apparent stability increases.

**Figure 8:**
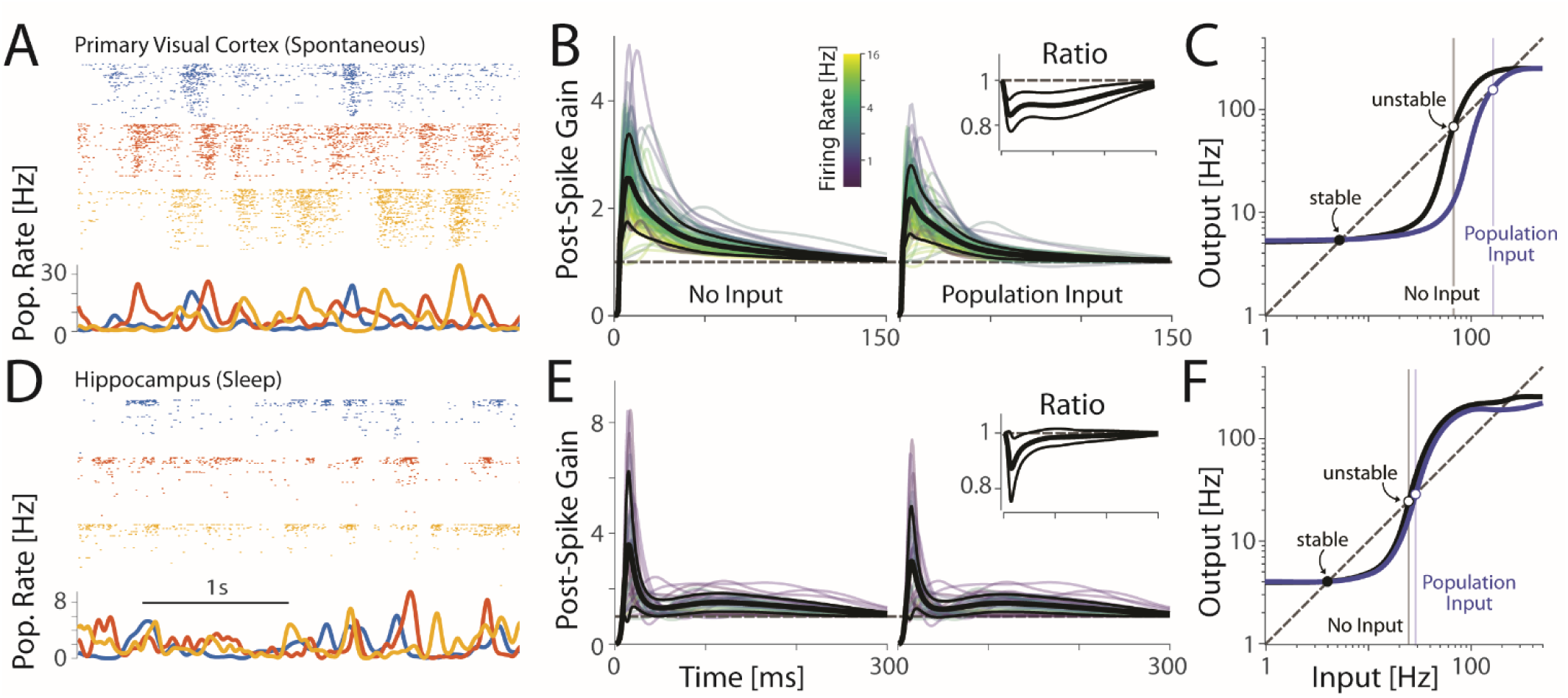
Post-spike filters estimated from real data decrease when population activity is included as a covariate. Segments of spontaneous activity are shown for V1 (A) and during sleep for hippocampus (D). Neurons are sorted by firing rate. B and E show estimated post-spike filters. Black lines denote the average filter (thick) and standard deviation (thin). For clarity, only filters for neurons with firing rates >1Hz are shown. C and F show average quasi-renewal transfer functions for the same set of neurons. All neurons appear to have fragile dynamics with one stable fixed point near the neuron’s average firing rate and an unstable fixed point, beyond which the neuron’s firing rate diverges. Including population covariates increases the region of stability.

## Discussion

When the goal of modeling is causal inference or understanding of biological mechanisms, the potential for biases due to omitted variables is often clear. The statistical effects of confounders (Wasserman, 2004), as well, as the limits that they place on neuroscientific understanding are widely appreciated (Jonas and Kording, 2017; Krakauer et al., 2017; Yoshihara and Yoshihara, 2018). However, when the goal of modeling is to create an abstract, explanation or summary of observed neural activity, the fact that omitted variables can bias these explanations is not always widely acknowledged. Here we have illustrated the potential for omitted variable bias in two types of commonly used GLMs for neural spiking activity: tuning curve models using spike counts across trials and models that capture single-neuron dynamics with a post-spike history filter. In each model, adding a previously omitted variable, as expected, improved spike prediction accuracy. However, what we emphasize here is that, when omitted variables were included, the estimates of the original parameters changed. For three case studies using tuning curves we found that by adding a traditionally omitted variable tuning curves showed less modulation due to the originally included variables. In models of single neuron dynamics, adding omitted variables led to decreased post-spike gain and greater apparent stability. Importantly, omitted variables can arise in GLMs in any situation where an omitted variable affects neural activity and the effect of the omitted variable is not independent of the included variables.

The case studies here are not unique, and many studies have described how adding additional variables to a tuning curve or single neuron model can improve prediction accuracy. In M1, in addition to movement speed, joint angles, muscle activity, end-point force, and many other variables also appear to modulate neural responses (Fetz, 1992; Kalaska, 2009). In addition to speed and head direction in the hippocampus, theta-band LFP, sharp-wave ripples, and environmental features, such as borders, appear to modulate neural activity (Hartley et al., 2014). And in V1, there is growing evidence that population activity (Lin et al., 2015) and non-visual information (Ghazanfar and Schroeder, 2006) modulates neural responses. In each of these systems, neural responses are affected by many, many factors. Responding to many task variables may even be functional, allowing downstream neurons to more effectively discriminate inputs (Fusi et al., 2016). In any case, it seems clear that our models do not yet capture the full complexity of neural responses (Carandini et al., 2005). By omitting relevant variables, current models are likely to be not just less accurate but also biased.

Parameter bias may be problematic in and of itself. However, omitted variable bias may also have an important effect on generalization performance. As noted in (Box, 1966), in a new context, the effect of the omitted variables and the relationship between the omitted and included variables may be different. Since the parameters of the included variables are biased, this change can reduce generalization accuracy. This phenomena may explain, to some extent, why tuning models fit in one condition often do not generalize to others (Graf et al., 2011; Oby et al., 2013). For models of single-neuron dynamics, omitted variable bias can also have a negative effect on the accuracy of simulations. Previous work has shown that simulating a GLM with post-spike filters estimated from data often results in unstable, diverging simulations. Although several methods for stabilizing these simulations have recently been developed (Gerhard et al., 2017; Hocker and Park, 2017), one, perhaps primary, reason for this instability may be that the post-spike filters are biased due to omitted synaptic input. Since estimated post-spike filters may reflect not just intrinsic neuron properties but also the statistics of the input, interpreting and comparing post-spike filters may be difficult. Different history parameters may be different due to intrinsic biophysics (Tripathy et al., 2013) or due to differing input, and resolving this ambiguity will likely involve more accurately accounting for the input itself (Kim and Shinomoto, 2012).

The possibility of omitted variable bias does not mean that estimated parameters, predictions, and simulations from simplified model are useless, but it may mean that we need to be cautious in interpreting these models and their outputs. When reporting the results of regression, in addition to avoiding describing associations with causal language, it may be generally useful to discuss known and potential confounds. Previous studies have already identified several specific cases of omitted variable bias where careful interpretation is necessary. For instance, omitted common input can bias estimates of interactions between neurons (Brody, 1999), and omitted history effects can bias receptive field estimates (Pillow and Simoncelli, 2003). In estimating peri-stimulus time histograms, omitting variables that account for trial-to-trial variation may cause biases (Czanner et al., 2008) or issues with identifiability (Amarasingham et al., 2015). Similarly, biases due to spike sorting errors (Ventura, 2009) could be framed as a result of omitting variables related to missing/excess spikes. Since we typically do not model or observe all the variables that affect neural activity, omitted variable problems are likely to be pervasive in systems neuroscience far beyond these specific cases.

Although we have focused on GLMs here, omitted variable bias can affect any model and other types of model misspecification can also result in biased parameter estimates. Adding input nonlinearities (Ahrens et al., 2008; David et al., 2009), interaction effects (McFarland et al., 2013), or higher-order terms to the GLM (Berger et al., 2010; Park et al., 2013) may fix certain types of model misspecification, but any model that omits relevant variables is still likely to suffer from the same problems. This includes both machine learning methods that may provide better prediction accuracy than GLMs (Benjamin et al., 2017) and single neuron models aiming to describe greater biophysical detail (Herz et al., 2006). Unlike over-fitting or non-convergence (Zhao and Iyengar, 2010), omitted variable bias will generally not be resolved by including additional data or by adding regularization. Moreover, adding one omitted variable, as we have done with the case studies here, is no guarantee that there are not other relevant variables being omitted.

One approach that could potentially reduce omitted variable bias is latent variable modeling, where the effects of unknown covariates are explicitly included (constrained by simplifying assumptions). Recent work has introduced latent variables for neural activity with linear dynamics (Smith and Brown, 2003; Kulkarni and Paninski, 2007; Paninski et al., 2010), switching dynamics (Putzky et al., 2014), rectification (Whiteway and Butts, 2017), and oscillations (Arai and Kass, 2017). And these models appear to out-perform GLMs on population data in retina (Vidne et al., 2012), visual (Archer et al., 2014), and motor cortices (Chase et al., 2010; Macke et al., 2011). Inferring latent variables requires making (sometimes strong) assumptions about the nature of the variables and may require observations from multiple neurons or across multiple trials, but, by approximating some of the effects of relevant omitted variables, latent variables may reduce omitted variable bias. However, generally determining when relevant variables are omitted from a model and what those variables are is not a trivial problem.

There is a well-known aphorism from George EP Box that, “All models are wrong, but some are useful.” The lengthier version of this quip is, “All models are approximations. Essentially, all models are wrong, but some are useful. However, the approximate nature of the model must always be borne in mind.” GLMs are certainly useful descriptions of neural activity. They are computationally tractable, can disentangle the relative influence of multiple covariates, and often provide the core components for Bayesian decoders. Here we emphasize, however, one ubiquitous circumstance in systems neuroscience where the “approximate nature” of the models should be “borne in mind.” Namely, omitted variables can bias estimates of the included effects.

## Methods

### Neural Data

All data analyzed here was previously recorded and shared by other researchers through the Collaborative Research in Computation Neuroscience (CRCNS) Data Sharing Initiative (crcns.org).

Data from primary motor cortex is from CRCNS dataset DREAM-Stevenson_2011 (Walker and Kording, 2013). These data were recorded using a 100-electrode Utah array (Blackrock Microsystems, 400 mm spacing, 1.5 mm length) chronically implanted in the arm area of primary motor cortex of an adult macaque monkey. The monkey made center-out reaches in a 20x20cm workspace while seated in a primate chair, grasping a two-link manipulandum in the horizontal plane (arm roughly in a sagittal plane). Each trial for the center-out task began with a hold period at a center target (0.3–0.5 s). After a go cue, subjects had 1.25 s to reach one of eight peripheral targets and then held this outer target for at least 0.2–0.5 s. Each success was rewarded with juice, and feedback (1-2cm diam) about arm position was displayed onscreen as a circular cursor. Spike sorting was performed by manual cluster cutting using an offline sorter (Plexon, Inc) with waveforms classified as single- or multi-unit based on action potential shape and minimum ISIs greater than 1.6 ms (yielding n=81 single units). Here we take model tuning curves using spike counts between 150ms before to 350ms after the speed reached its half-max. Average movement speed for each trial was calculated from 0-250ms after the speed reached its half-max (290 trials in total). For the details of the surgery, recording, and spike sorting see (Stevenson et al., 2011).

Hippocampal data is from CRCNS HC-3 (Mizuseki et al., 2013). Here we use recording sessions ec16.19.272 and ec014.215, where a Long-Evans rat was sleeping and foraging in a 180x180cm maze, respectively. For both recordings, 12-shank silicon probes (with 8 recording sites each, 20μm separation) were implanted in CA1 (8 shanks) and EC3-5 (4 shanks) (based on histology). Spikes were sorted automatically using KlustaKwick and then manually adjusted (Klusters) yielding 85 units for the sleep data and 117 for the open field data. For the sleep data where we model post-spike history effects, the spike trains were binned at 1ms and the recording length was 27min. Here we model all neurons with firing rates >0.5Hz (n=39). For the open field data where we model place tuning, spike trains were binned at 250ms and the recording length was 93min. Place cells (n=68) were selected based on having an overall firing rate <5Hz (to rule out interneurons), a peak firing rate >2Hz, and a contiguous set of pixels (after smoothing with an isotropic Gaussian *σ*=8cm) of at least 200 cm^2^ where the firing rate was above 10% of the peak rate. For details of the surgery, recording, and spike sorting see (Diba and Buzsaki, 2008; Mizuseki et al., 2009).

Data from primary visual cortex is from CRCNS dataset PVC-11 (Kohn and Smith, 2016). Here we use spontaneous activity, during a gray screen, (Monkey 1) and responses to drifting sine-wave gratings (Monkey 1) both from an anesthetized (Sufentanil - 4-18 microg/kg/hr) adult monkey (Macaca fascicularis). Recordings were made in parafoveal V1 (RFs within 5 degrees of the fovea) using a 96-channel multi-electrode array (Blackrock Microsystems), 400 mm electrode spacing, 1mm depth. After automatic spike sorting and manual cluster adjustment, 87 and 106 units were recorded during spontaneous activity and grating presentation, respectively. Only neurons with waveform SNR>2 and firing rates >1Hz were analyzed, n=62 for spontaneous and n=90 for grating data. For the spontaneous activity we bin spike counts at 1ms and the recording length was 20min. For the drifting grating data, we analyzed spike counts from 200ms to 1.2s after stimulus onset on each trial – 12 directions, 2400 trials total. Gratings had a spatial frequency of 1.3 cyc/deg, temporal frequency of 6.25Hz, size of 8-10 deg (to cover receptive fields of all recorded neurons) and were presented for 1.25s with a 1.5s inter-trial interval between stimuli. For surgical, stimulus, and preprocessing details see (Smith and Kohn, 2008; Kelly et al., 2010).

### Tuning Curve Models

For the M1 data we use a circular, cubic B-spline basis with 5 equally spaced knots

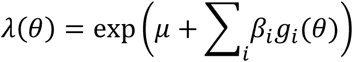

where *g*(·) are the splines that depend on the reach direction *θ*, weighted by parameters *β* and the parameter *μ* defines a baseline firing rate. To include the effect of speed, we then add three covariates

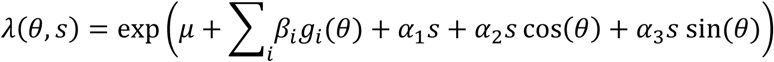

where *s* indicates the speed, and the parameters *α* allow for a multiplicative speed effect as well as possible cosine-tuned speed x direction interactions as in (Moran and Schwartz, 1999).

For place fields in hippocampus we use isotropic Gaussian radial basis functions *f*(·) equally spaced (30cm) on an 6x6 square lattice with a standard deviation of 30cm

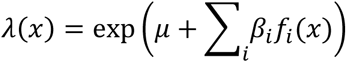

We find that the effect of speed is well modeled using the log-transformed speed *s*, and to model head direction-dependence we use circular, cubic B-splines *g*(·) with 6 equally spaced knots

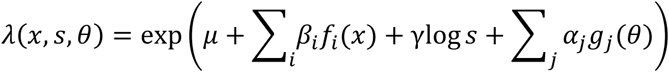

For the V1 data we again use a circular, cubic B-spline basis for the direction of the sine-wave grating (7 equally spaced knots).

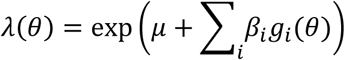

We find that the effect of population activity is well modeled using the total log-transformed firing rate of all neuron’s excepting the one being modeled

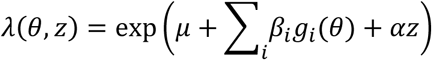

where *z* = ∑_*i*≠*j*_ log(*n*_*i*_ + 1). In all models, to avoid overfitting, especially for low firing rate neurons, we add a small L2 penalty to the log-likelihood with a fixed hyperparameter of 10^-4^.

### Post-spike History Simulations and Population Rate Models

In addition to capturing tuning curves, many studies have used GLMs to describe the dynamics of single spike trains (Brillinger, 1988; Harris et al., 2003; Paninski, 2004; Okatan et al., 2005; Truccolo et al., 2005; Weber and Pillow, 2017). Here, to account for post-spike history effects, we use a GLM taking the form

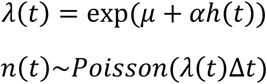

where *h*(*t*) denotes the vector of spike history covariates representing the recent history of spiking and *μ* determines a baseline firing rate. Here we assume *h*_*i*_(*t*) = ∑_τ>0_ *f*_*i*_(τ)*n*(*t* – τ), and we use neuron-specific, cubic B-spline bases *f*(·) whose knots are determined by the quantiles of each neuron’s ISI distribution. Specifically, we choose knots spaced between 10 and 400ms (HC) or 2 and 200ms (V1), where the spacing follows equal percentile regions of the ISI distribution in that same range. This gives 6 basis functions, and coefficients *α* to capture the spike-history. To enforce refractoriness, we fix the coefficient of the fastest basis (which peaks at 0 and ends at 10ms) to be -5, leaving 5 coefficients to be estimated.

The population rate model simply adds covariates where, for each neuron *i*

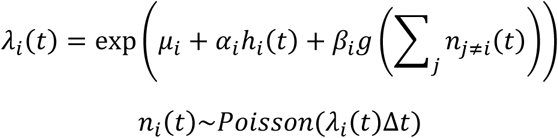

Here we use a set of acausal Gaussian filters for *g*(·) with standard deviations 20, 50, and 100ms. Note that spikes from the neuron being modeled are excluded from the population covariates.

### Stability analysis

Here we make use of a stability analysis proposed in (Gerhard et al., 2017). Briefly, we use a quasi-renewal approximation of the conditional intensity by considering the effect of the most recent spike, at time *t*′, and averaging over possible spike histories preceding this spike

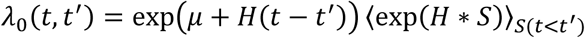

where *H*(*t* – *t*′) = *αf*(*t* – *t*′) and *S* represents the history of spiking. By assuming that *S* is generated from a homogeneous Poisson process with firing rate *A*_0_, the second term can be approximated by

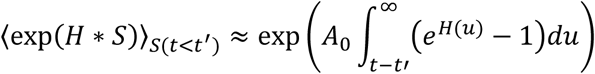

Given this approximation, we can then estimate the inter-spike interval distribution as we would for a true renewal process and the steady-state distribution of inter-spike intervals is given by

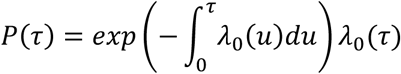

and the predicted steady-state firing rate is *f*(*A*_0_) = 1*/E*_*P*(τ)_[τ].

To assess stability, we can then examine how the predicted steady-state firing rate depends on the assumed rate of the homogeneous Poisson process *A*_0_. In particular, when *f*(*A*_0_) = *A*_0_ the quasi-renewal model has a fixed-point. To allow for external input, we incorporate the average effect of the covariates *X* into the conditional intensity approximation

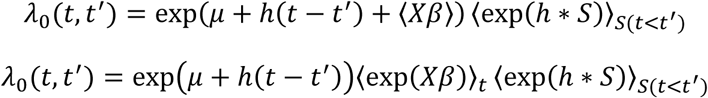

Note that, in general, adding inputs *X* will only change the stability of the model to the extent that these covariates change the estimate of *h*.

